# Naïve chicks prefer hollow objects

**DOI:** 10.1101/050799

**Authors:** Elisabetta Versace, Jana Schill, Andrea Maria Nencini, Giorgio Vallortigara

**Affiliations:** Center for Mind/Brain Sciences, University of Trento (Italy); Universität Osnabrück, Germany

**Keywords:** Predispositions, hollow, filled, innards, *Gallus gallus*, filial imprinting, chicks

## Abstract

Biological predispositions influence approach and avoid responses since the beginning of life. Neonates of species that require parental care (e.g. human babies and chicks of the domestic fowl) are attracted by stimuli associated with animate social partners, such as face-like configurations, biological motion and self-propulsion. The property of being filled is used as a cue of animacy by 8-month-old infants but it is not known whether this reflects the effect of previous experience. We use chicks of the domestic fowl (*Gallus gallus*) to investigate whether the property of being filled *vs.* hollow elicits spontaneous or learned preferences. To this aim we tested preferences of naΪve and imprinted chicks for hollow and closed cylinders. Contrary to our expectations, we documented an unlearned attraction for hollow stimuli. The preference for hollow stimuli decreased when chicks were imprinted on filled stimuli but did not increase when chicks were imprinted on hollow stimuli, suggesting that this feature is not crucial to categorize the familiarity of imprinting objects. When chicks were imprinted on occluded stimuli that could be either filled or hollow, the preference for hollow stimuli emerged again, showing that imprinting does not disrupt the spontaneous preference for hollow objects. Further experiments revealed that hollow objects were mainly attractive by means of depth cues such as darker innards, more than as places to hide or as objects with high contrast. Our findings point to predisposed preferences for hollow objects that might be unrelated to social behaviour.

## Introduction

Sensory and cognitive predispositions influence approach and avoid responses since the beginning of life [1–3]. In different species we observe spontaneous preferences for specific colours [4–7], shapes and sizes [6,8,9], configurations [10], dynamics [11,12], and odours [13–16]. In precocial species, individuals are mobile soon after birth, and can be tested when they have little if any experience, to investigate spontaneous preferences [3]. Soon after hatching, chicks of the domestic fowl (*Gallus gallus*), which is a nidifugal species, possess some spontaneous preferences to approach stimuli that are associated with animate social partners [17]. When given a choice between a stuffed hen and a stuffed scrambled hen, chicks prefer to approach the hen [18,19]. The same preference is consistent across different breeds [20]. Behavioural studies have found that this preference is driven by an unlearned attraction towards the face configuration contained in the stuffed hen [10,18]. Moreover, between the biological movement of a hen or a cat and the rigid motion of a hen rotated on its vertical axis, chicks prefer to approach the biologically moving object [11,21]; and between a self-propelled object and an object propelled by another one, naïve chicks prefer the self-propelled object [12]. Overall, chicks prefer to approach objects which are endowed with more animate features [2,3,22].

Observations on infants [23] suggest that 3-year-old children have a representation of the insides of animate beings as more likely to be filled than those of inanimate objects. Studies on human infants [24] have shown that 8-month-old babies possess expectations about the biological properties of animate and agentive entities. In this study infants were more surprised to see that self-propelled and agentive objects were hollow than when there was no evidence that those objects were hollow. It is not clear though whether previous experience with animate entities with innards (e.g. the parents) had generated infants' expectations, or whether they arose spontaneously. We reasoned that chicks of the domestic fowl (*Gallus gallus*), which are spontaneously attracted by entities which show cues associated with animacy in the absence of previous experience [2,3], might be a convenient subject to identify whether the property of being filled/hollow triggers unlearned preferences.

To this aim we tested preferences of naïve chicks (**Experiment 1**) maintained in darkness (Experiment 1a) or exposed to light (Experiment 1b) for hollow and closed cylinders of the size and colour that elicit filial responses. Moreover, since chicks rapidly learn features of their social partners by mere exposure through filial imprinting [25,26], they are a valuable model to study the role of experience in modifying spontaneous preferences. To this aim we investigated how imprinting modified unlearned preferences for hollow and filled objects (**Experiment 2**) after imprinting on hollow objects (Experiment 2a), filled objects (Experiment 2b) and objects who could not be perceived hollow or filled because their sides were occluded (Experiment 2c).

Since we noticed an overall preference for hollow objects, we investigated whether this behaviour was elicited by a preference for the stimulus that could better hide the chick (chicks could enter the hollow stimulus). In **Experiment 3** we checked whether the preference for hollow stimuli was still present when the stimuli were too small to host and hide chicks. We tested both dark-reared chicks and chicks exposed to light that had never seen the test stimuli or any other object of similar size, shape and colour. We observed a preference for hollow objects. In **Experiment 4** we checked whether the size of the hollow object was important in determining the preference for hollow objects comparing the preference for the large and the narrow hollow objects. In **Experiment 5** we checked whether the darker colour of the shadows present in the innards of hollow objects has a role in driving preferences for hollow stimuli by comparing preferences for filled objects with a white *vs.* a black stopper (Experiment 5a). Since chicks preferred the object with the black stopper, we tested whether the preference for a hollow stimulus was stronger or weaker than the preference for a black cap (Experiment 5b). The observed preference for the black cap stimulus could be explained both by brightness (chicks preferred lower brightness) and by contrast (chicks preferred greater contrast). To clarify the importance of contrast and brightness in determining the preference for hollow objects, in **Experiment 6** we used two-dimensional stimuli with different colour and identical contrast, i.e. a white disk on a black background *vs.* a black disk on a white background. If the preference of chicks for hollow *vs.* Filled and for Black *vs.* Hollow was driven by the darker colour (innards or cap), in this contrast chicks should have chosen the white disk on a black background. If the preference was driven by contrast, chicks were expected to have no preference. A preference for the black disk on a white background would be consistent with a preference for darker objects/innards, possibly a cue of depth.

## MATERIALS AND METHODS

### Ethical note

All applicable international, national, and/or institutional guidelines for the care and use of animals were followed. This study was approved by the ethical committee of the University of Trento (Organismo Preposto al Benessere degli Animali) prot. N. 14-2015 and was licensed by the Ministero della Salute, authorization n. 1138/2015. The research adhered to the ASAB/ABS Guidelines for the Use of Animals in Research.

### Subjects

The subjects were 24-hour old chicks of the domestic fowl (*Gallus gallus*) of the Hybro strain (a local hybrid variety of the White Leghorn breed). This breed has been selected to be sexually dimorphic at the moment of hatching, and chicks can be easily sexed looking at their feathers. The eggs were obtained from a commercial hatchery (Agricola Berica, Montegalda, Italy), then incubated in complete darkness (as in other experiments on predispositions) at 37.7 °C until hatching, with the same procedure used in other tests for spontaneous preferences in chicks [11,12,19]. Three days before hatching humidity was increased from 40% to 60%. Eggs hatched in individual boxes (11 × 8.5 × 14 cm) and chicks could hear their conspecifics but had no visual or tactile contact with conspecifics until the moment of test. The exact number of chicks used in each experiment, divided by Sex and presence/absence of choice during the test, is presented in Table 1 (chicks that did not move from the central area were excluded from the analyses since did not show any preference).

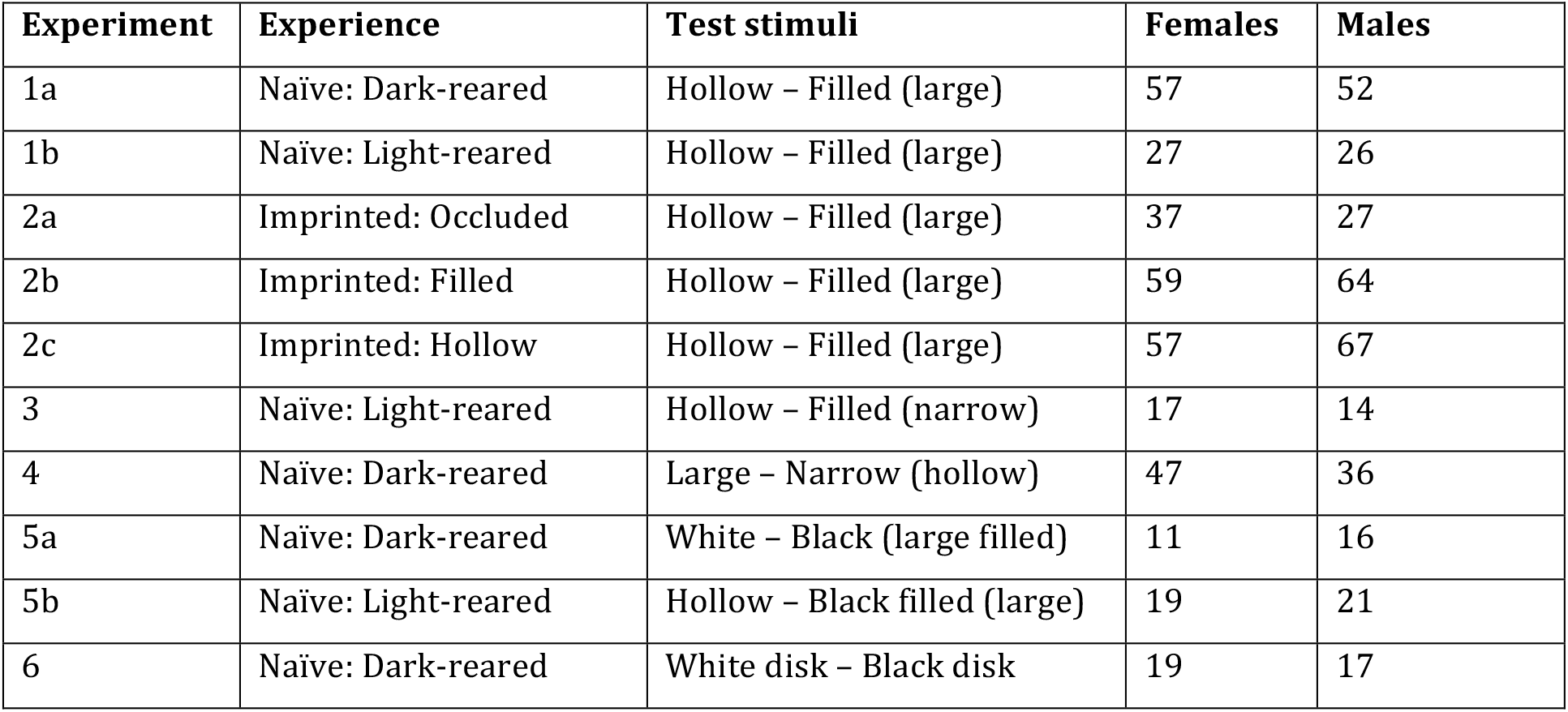

### Test stimuli

Test stimuli are shown in Figure 1. In Experiment 1 and 2 test stimuli were large plastic tubes (12 cm, ø 4 cm) left open (Hollow, Fig. 1A) or closed with a white cap (Filled, Fig. 1B), with an orange external surface and a white internal surface. In Experiment 3 we used the same stimuli with the only difference that the diameter was 2.5 cm (Narrow stimuli are shown in Fig. 1C and 1D). In Experiment 4 we used Large and Narrow hollow stimuli (Fig. 1A and 1C). In Experiment 5 we used stimuli similar to those used in Experiments 1 and 2 with the only difference that one cap was black (Fig. 1E). In Experiment 6 we used a white disk on a black background, and a black disk on a white background (Fig. 1F) with a diameter of 4 cm located at 4.5 cm from the ground.

**Figure 1.**
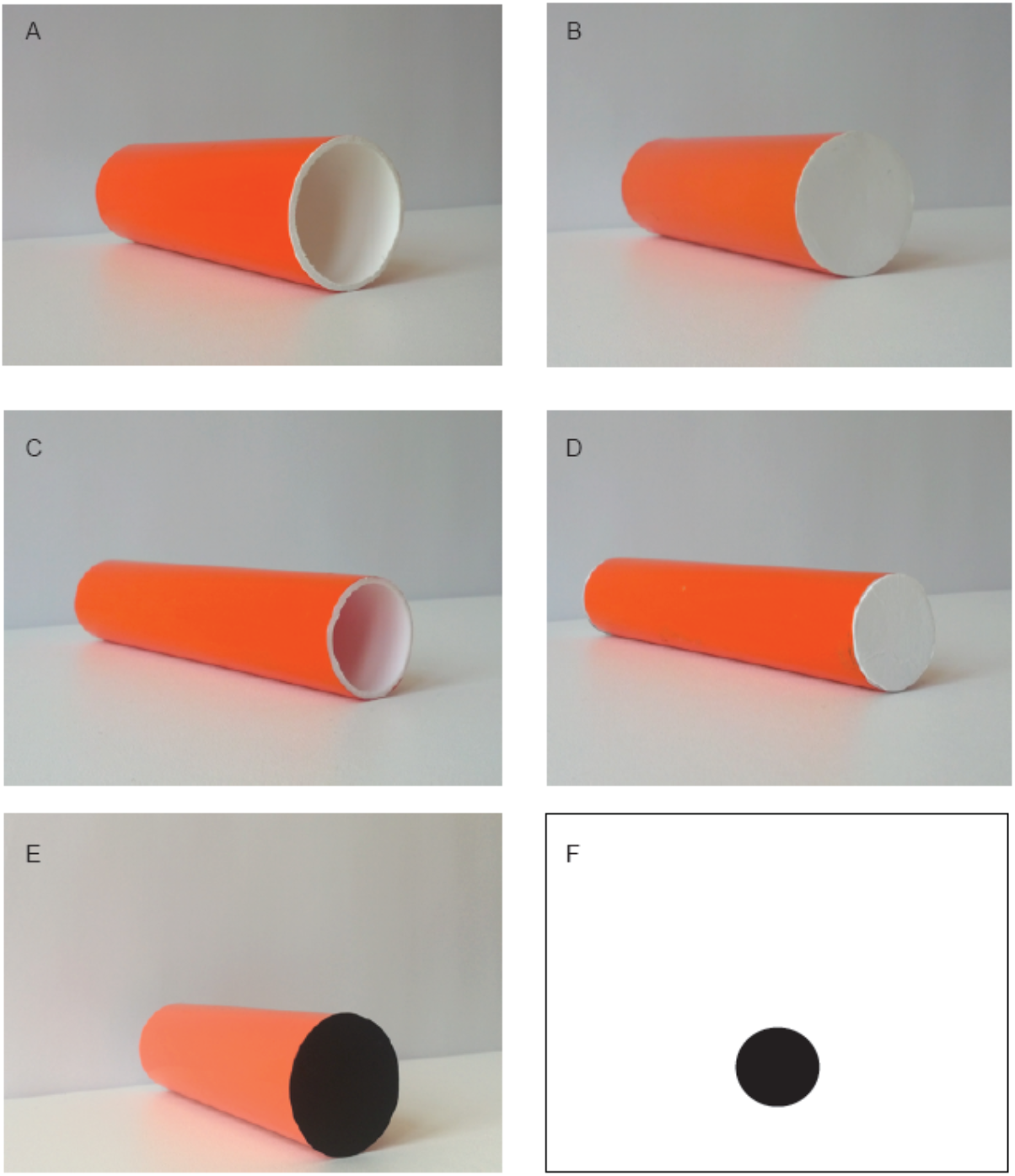
Stimuli used in Experiment 1 and 2 (**A** and **B**), Experiment 3 (**C** and **D**), Experiment 4 (**A** and **C**), Experiment 5 (**A** and **E**), and Experiment 6 (Panel **F** shows the Black disk on the white background. The other stimulus was a White disk on a black background).

### Imprinting stimuli

In experiment 2 chicks were individually imprinted to orange cylinders (12 cm, ø 4 cm), that were presented through a 7.5 × 10 cm transparent plastic window. Imprinting lasted 24 ± 3 and was immediately followed by the test. Chicks had no direct interaction with the stimulus during imprinting and the only interaction with conspecifics was auditory. In the Occluded condition the cylinder was presented horizontally and the chicks could not see whether it was hollow or filled because the edges were covered. In the Hollow and Filled condition the hollow and or filled cylinder were presented perpendicular to the transparent window and the chick could see whether it was hollow or not.

### Test apparatus

The experiment took place in a 100 × 30 × 31 cm white arena open on the top (see Figure 2). Test stimuli were located in the middle of each short side on a white plastic platform that was 4.5 cm high. The box was virtually divided into three areas: a left area (41 cm), a central area (18 cm) and a right area (41 cm). The white platforms occupied 15 cm in each side area. In Experiment 4, the platforms were removed and the stimuli were placed directly on the walls of the apparatus. The right-left position of the stimuli was counterbalance between subjects.

**Figure 2.**
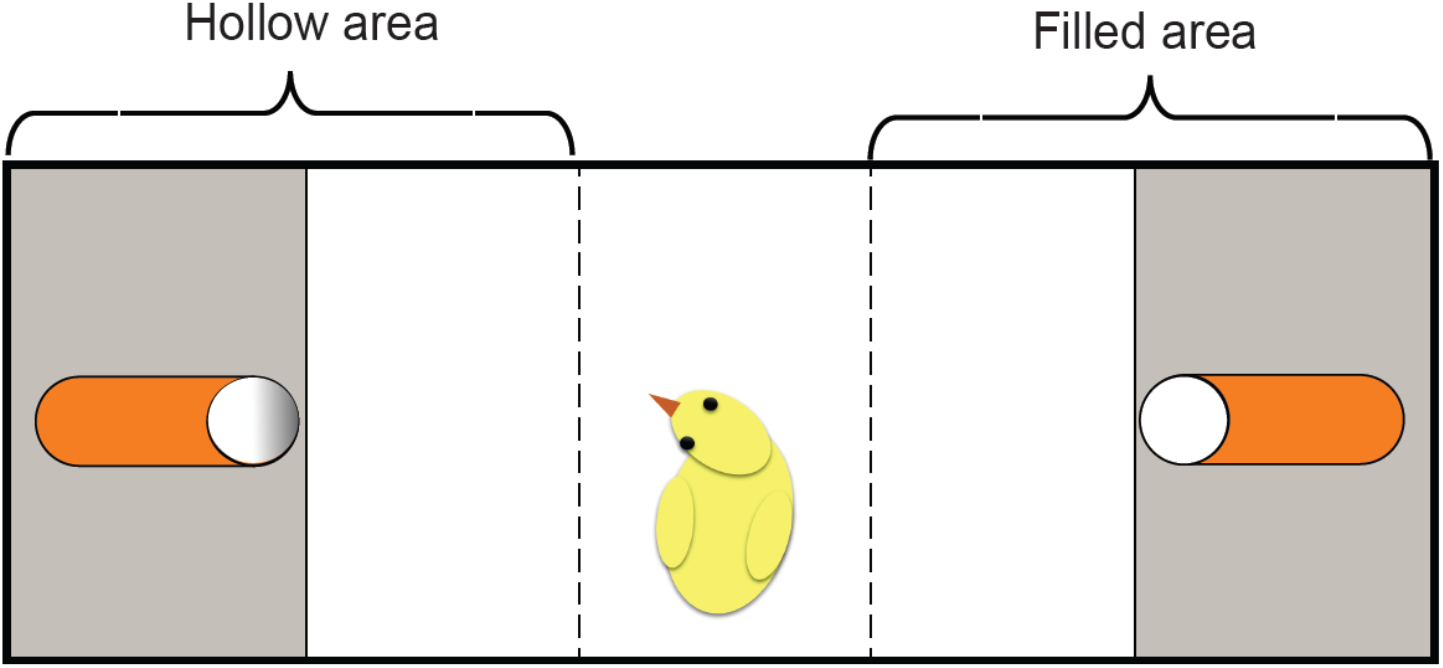
Illustration of the testing apparatus. The right/left position of the stimuli was counterbalanced between subjects.

## PROCEDURE

### Imprinting

Soon after hatching, in the imprinting experiments chicks were individually exposed to the imprinting stimulus for 24 hours before testing under constant light. Imprinting cages were 28 × 38 × 32 cm and the stimulus was presented through a transparent partition (7.5 × 10 cm). In this way chicks had no direct interaction with the stimuli before testing, similarly to naïve chicks that had never experienced stimuli like those used during the test.

### Test

#### Procedure and data analysis

We followed the same procedure in all experiments. Each chick was individually located in the centre area facing the long side of the box opposite to the experimenter and video recorded for 360 seconds. We recorded which side area was entered first (First choice) and the seconds spent in each side area. The chick was considered to have entered a new sector as soon as it crossed the borderline with both feet. After the testing phase chicks were not used in any other experiment. For the chicks which entered side areas that indicate a choice we checked whether the first choice was significantly different from the 0.5 chance level using a Chi-squared test, with alpha = 0.05.

For each chick that left the central area we calculated an index of preference for the Hollow stimulus (Experiments 1, 2 and 3) or an index of preference for the Narrow (Experiment 4) or Black stimulus (5) in this way:

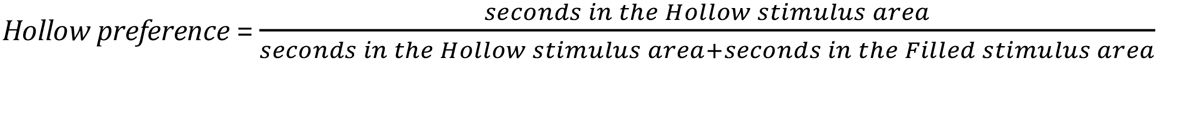

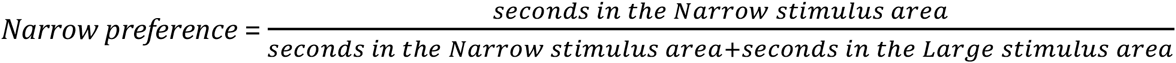

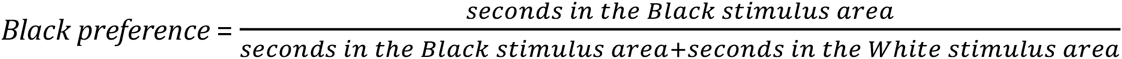

For all indices, 1 indicates a full preference for the respective stimulus (Hollow, Narrow, Black), 0.5 no preference and 0 a full preference for the opposite stimulus (Filled, Large, White). Since all data had a bimodal distribution with peaks on the extremes (0 and 1) we used non-parametric statistics to test for significance: the Kruskal-Wallis test to test for differences between conditions and sexes, and the Mann-Whitney-Wilcoxon one-sample test *vs.* the 0.5 chance level.

Chicks that did not make any choice were excluded from the analyses on the preferences but we compared the chicks that made a choice between naïve and imprinting experiment to check for the effectiveness of the imprinting procedure: we expected a higher choice rate in imprinted chicks compared to naïve chicks.

## RESULTS

### Experiment 1: naïve chicks (dark-reared and light-reared) chicks tested with Hollow *vs.* Filled stimuli

We assessed the preference for the hollow/filled object in naïve chicks, namely dark-reared and light-reared chicks that had never experienced any of the test stimuli before the test.

#### First choice

There was no significant difference between dark- and light-reared chicks (Chisquare test: *χ* = 0.073, df = 1, *P* = 0.79), and in both conditions chicks had the same trend, therefore we collapsed the two naïve conditions for further analyses. The number of chicks that approached the Hollow *vs.* Filled stimulus was significantly different from chance (Chi-square test: *χ* = 8.91, df = 1, *P* = 0.003) with an overall preference for the Hollow stimulus (Figure 3A). Compared to dark-reared chicks, naïve chicks exposed to light or other stimulation are known to exhibit stronger predisposed preferences [11,27], and we could use a smaller sample for light-reared chicks.

**Figure 3.**
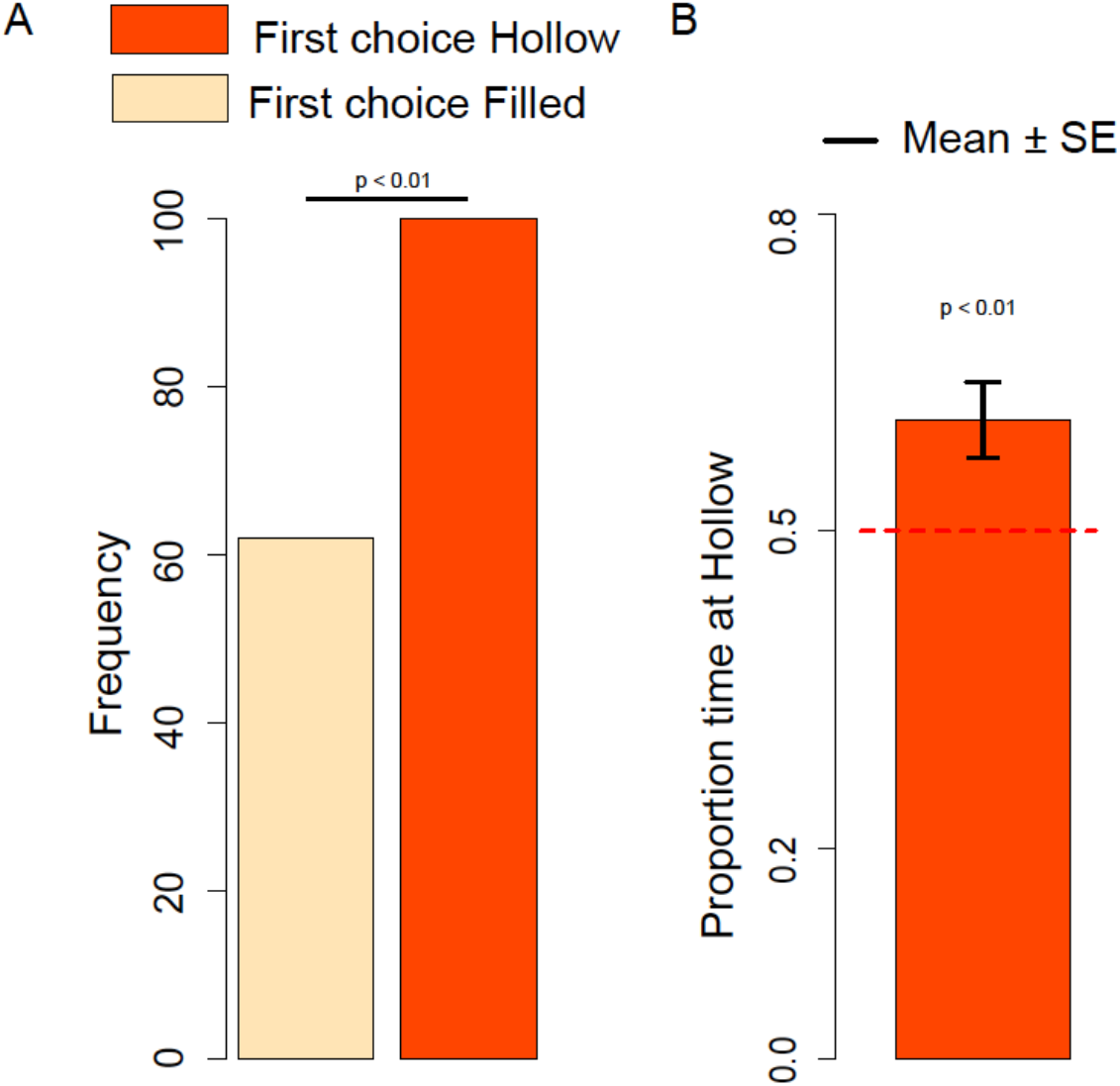
**A.** Number of naïve chicks that first approached the Hollow or Filled stimulus in the dark-reared and light-reared condition. **B.** Proportion of time spent at the Hollow stimulus by dark-reared and light-reared chicks.

#### Hollow preference

Considering the Hollow preference index we did not observe any significant Sex difference (Kruskal-Wallis test: *H* = 0.32, df = 1, *P* = 0.57) or Exposure (dark- *vs.* light-rearing) difference (Kruskal-Wallis test: *H* = 0.07, df = 1, *P* = 0.79), therefore we collapsed the two naïve conditions for further analyses. We documented a significant preference for the Hollow stimulus (Mann-Whitney test: *V* = 8053, df = 1, *P* = 0.01), see Figure 3B.

### Experiment 2: chicks imprinted with Hollow, Filled or Occluded and tested with Hollow *vs.* Filled stimuli

To investigate the role of experience in determining the preferences for hollow objects we investigated the preference for the hollow/filled object in imprinted chicks, namely chicks that had been exposed to the filled or hollow object, or to an object located horizontally the sides of which were occluded, so that it did not show whether it was filled or hollow.

#### First choice

The number of chicks that approached the Hollow *vs.* Filled stimulus was significantly different between imprinting conditions (Chi-square test: *χ* = 7.15, df = 2, *P* = 0.028). Chicks imprinted on the Occluded object showed a significant preference for the hollow object (Chi-square test: *χ* = 7.56, df = 1, *P* = 0.006), whereas chicks imprinted on the Filled (Chi-square test: *χ* = 0.40, df = 1, *P* = 0.53) and Hollow object (Chi-square test: *χ* = 2.61, df = 1, *P* = 0.11) did not. While the first choice of chicks imprinted on the Occluded object did not differ from the first choice of chicks imprinted on the Hollow object (Chi-square test: *χ* = 1.35, df = 1, *P* = 0.24), there was a significant difference between the first choice of chicks imprinted on the Occluded object and the first choice of chicks imprinted on the Filled object (Chi-square test: *χ* = 6.02, df = 1, *P* = 0.014). Only chicks imprinted on the Filled object had a tendency to choose the Filled object (Figure 4A). While running the experiments, we noticed a trend for a sex difference between hollow/filled imprinted chicks. In the light of the documented sex differences in the preference for the slight novelty of imprinting objects between male and female chicks [28,29], we decided to increase the sample in these conditions to clarify whether it was a spurious effect. After increasing the sample, the trend disappeared, but we ended up with a larger sample size for these two groups.

**Figure 4.**
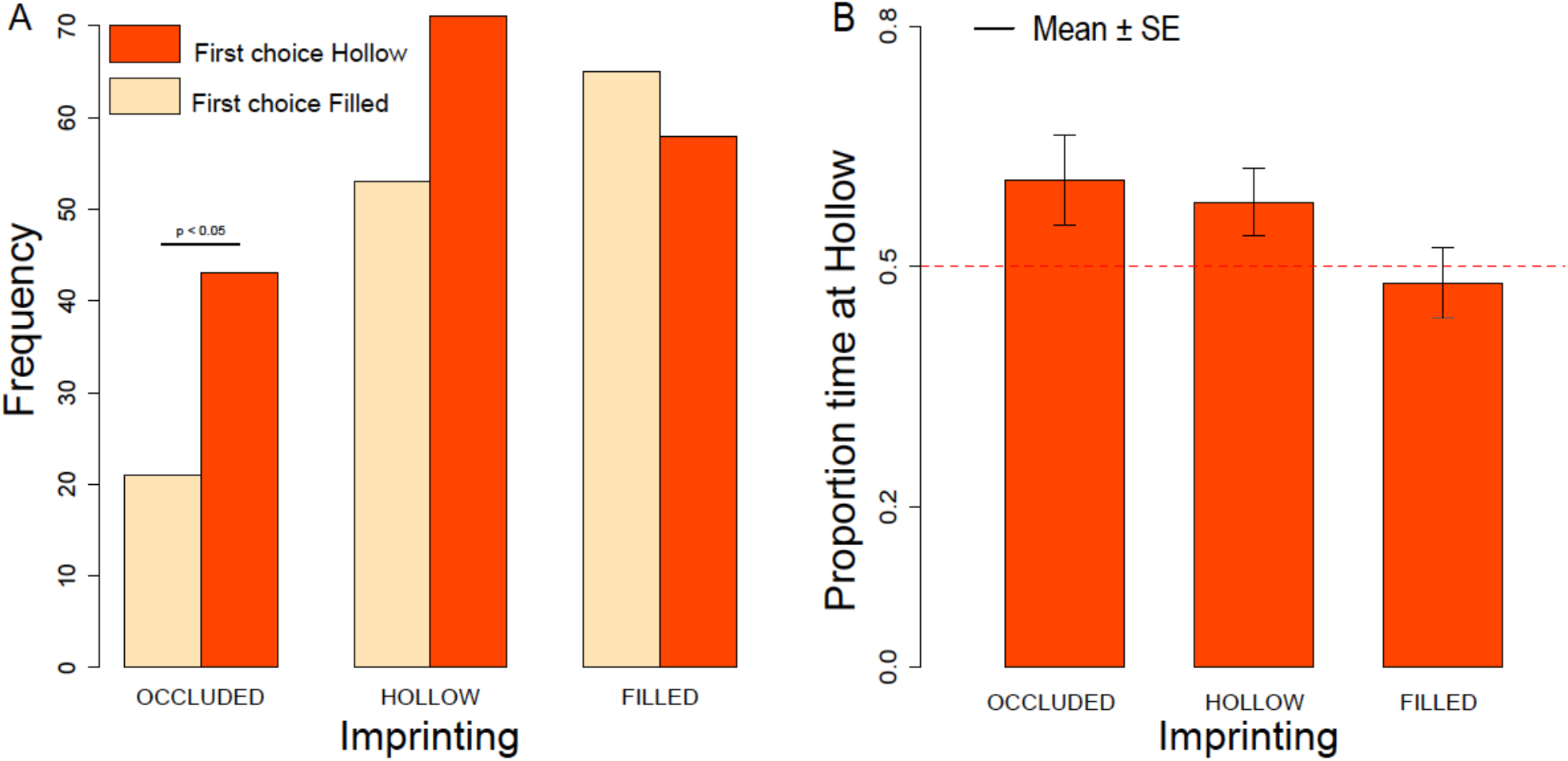
**A.** Number of imprinted chicks that first approached the Hollow or Filled stimulus after being exposed to Occluded, Hollow or Filled imprinting stimuli. **B.** Proportion of time spent at the Hollow stimulus for chicks exposed to Occluded, Hollow or Filled imprinting stimuli.

#### Hollow preference

Considering the Hollow preference index we did not observe any significant Sex difference (Kruskal-Wallis test: *H* = 1.60, df = 1, *P* = 0.21) or Exposure difference (Kruskal-Wallis test: *H* = 3.66, df = 2, *P* = 0.161). We observed an overall trend for preferring the Hollow stimulus (Mann-Whitney test: *V* = 27016.5, df = 1, *P* = 0.063), that turned out highly significant when considering only the chicks never exposed to filled stimuli, namely chicks imprinted on the occluded and hollow objects (Mann-Whitney test: *V* = 10721.5, df = 1, *P* = 0.009), see Figure 4B.

### Experiment 3: naïve chicks tested with narrow Hollow *vs.* narrow Filled stimuli

To investigate the extent and consistency of the hollow preference, we tested the preference for the hollow/filled object in naïve chicks, using smaller stimuli than those used in Experiment 1.

#### First choice

Chicks confirmed the preference for hollow stimuli (Chi-square test: *χ* = 17.06, df = 1, *P*< 0.001).

#### Hollow preference

Considering the Hollow preference index we did not observe any significant Sex difference (Kruskal-Wallis test: *H* = 1.46, df = 1, *P* = 0.23) but an overall preference for the Hollow stimulus (Mann-Whitney test: *V* = 461, df = 1, *P*< 0.001).

### Experiment 4: naïve chicks tested with Large hollow *vs.* Narrow hollow stimuli

To investigate whether the preference of young chicks for hollow objects was driven by the possibility to hide inside hollow objects, we presented naïve dark-reared chicks with a choice between Large (4 cm in diameter, large enough to hide a chick) and Narrow hollow stimuli (2.5 cm in diameter, too small to hide a chick).

#### First choice

The number of chicks that approached the Large *vs.* Narrow stimulus was not significantly different between Sexes (Chi-square test: *χ* = 0.14, df = 1, *P* = 0.71), therefore we collapsed the data from males and females together. There was no significant preference for the Large or Narrow stimulus (Chi-square test: *χ* = 0.108, df = 1, *P* = 0.74), suggesting that the possibility to hide inside the Large hollow stimuli is not the main drive of the preference for hollow stimuli.

#### Narrow preference

Considering the Narrow preference index we did not observe any significant Sex difference (Kruskal-Wallis test: *H* = 0.11, df = 1, *P* = 0.74). Overall we observed no significant preference for Large or narrow stimuli (Mann-Whitney test: *V* = 1583.5, df = 1, *P* = 0.56).

### Experiment 5a: naïve chicks tested with filled White *vs.* filled Black stimuli

#### First choice

The number of chicks that approached the White *vs.* Black stimulus was not significantly different between Sexes (Chi-squared test: *χ* = 0.12, df = 1, *P* = 0.73), therefore we collapsed the data from males and females together. We observed a significant preference for the Black stimulus (Chi-squared test: *χ* = 16.33, df = 1, *P* < 0.001), see Figure 5A.

**Figure 5.**
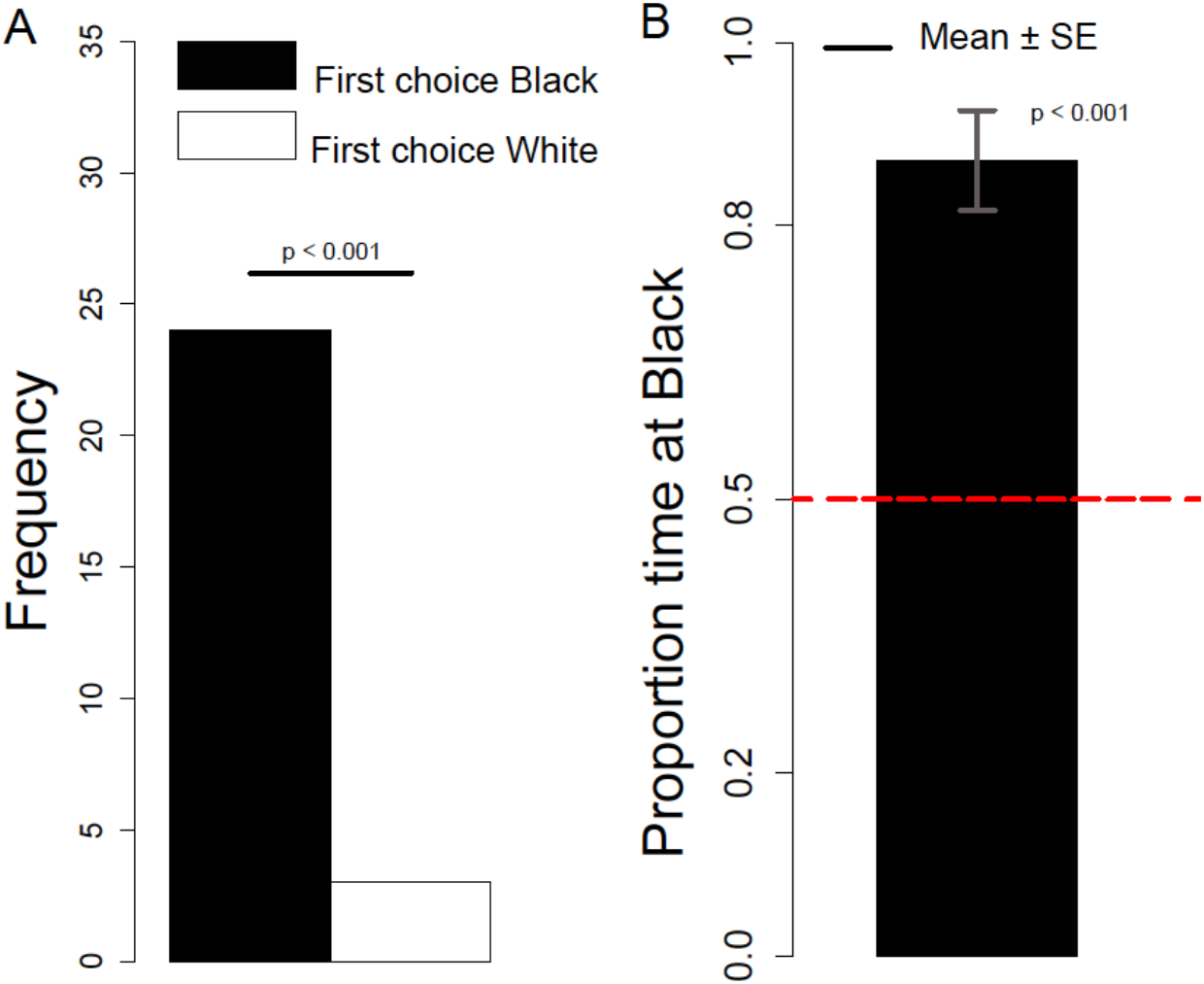
**A.** Number of chicks that first approached the Black or White stimulus. **B.** Proportion of time spent at the Black stimulus.

#### Black preference

Considering the Black preference index, we did not observe any significant Sex difference (Kruskal-Wallis test: *H* = 0.066, df = 1, *P* = 0.80). Overall we observed a significant preference for the Black stimulus (Mann-Whitney test: *V* = 354, df = 1, *P* < 0.001), see Figure 5B.

### Experiment 5b: naïve chicks tested with filled Black *vs.* Hollow stimuli

#### First choice

The number of chicks that approached the Hollow *vs.* Black stimulus was not significantly different between Sexes (Chi-squared test: *χ* = 0.307, df = 1, *P* = 0.58), therefore we collapsed the data from males and females together. We observed a significant preference for the Black stimulus (Chi-squared test: *χ* = 14.4, df = 1, *P* < 0.001), see Figure 6A.

**Figure 6.**
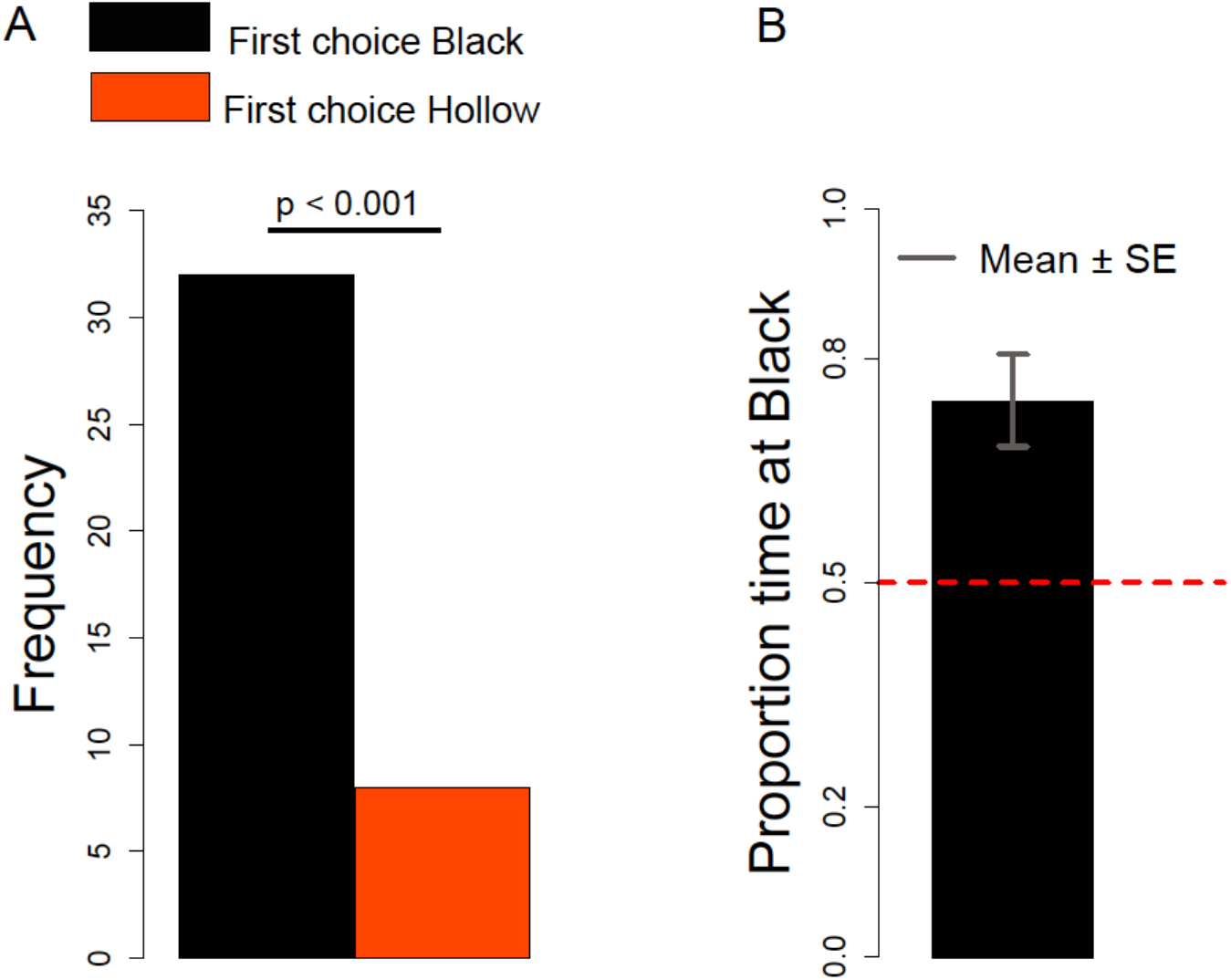
**A.** Number of chicks that first approached the Black or Hollow stimulus. **B.** Proportion of time spent at the Black stimulus.

#### Black preference

Considering the Black preference index we did not observe any significant Sex difference (Chi-squared test: *χ* = 0.818, df = 1, *P* = 0.366). Overall we observed a significant preference for the Black stimulus (Mann-Whitney test: *V* = 174, *P* < 0.001), see Figure 6B.

### Experiment 6: naïve chicks tested with a White disk on a black background *vs.* a Black disk on a white background

#### First choice

The number of chicks that approached the White *vs.* Black disk was not significantly different between Sexes (Chi-squared test: *χ* = 0.166, df = 1, *P* = 0.68). Overall we observed a significant preference for the Black stimulus (Chi-squared test: *χ* = 19.882, df = 1, *P* < 0.001), see Figure 7A.

**Figure 7.**
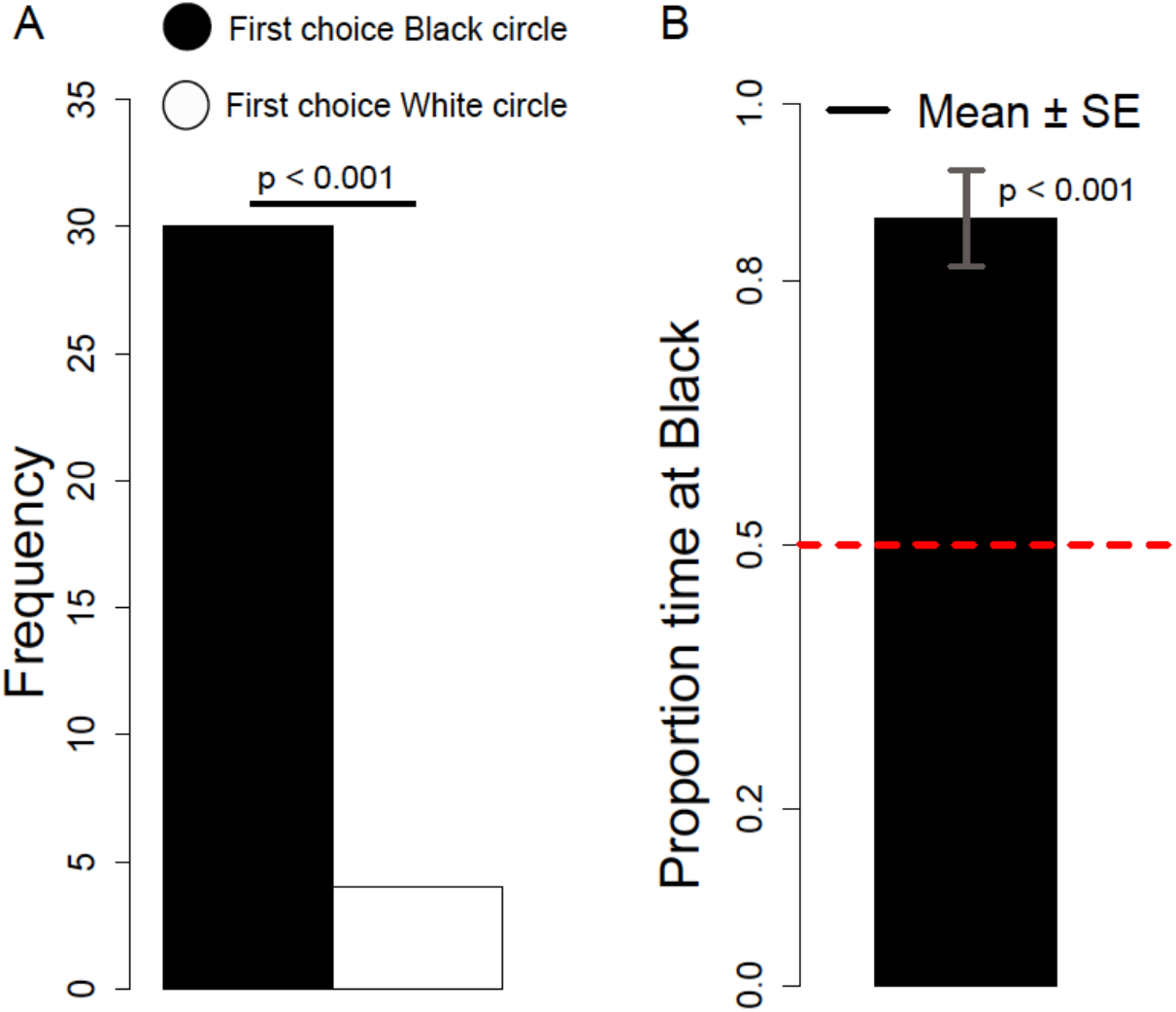
**A.** Number of chicks that first approached the Black disk on a white background or the White disk on a black background. **B.** Proportion of time spent at the Black disk on a white background.

#### Black preference

Considering the Black preference index we did not observe any significant Sex difference (Kruskal-Wallis *χ*^2^ = 0.65, df = 1, *P* = 0.42). Overall we observed a significant preference for the Black stimulus (Mann-Whitney test: *V* = 519, df = 1, *P* < 0.001), see Figure 7B.

## Discussion

Sensory and cognitive predispositions can help naïve individuals in deciding whether to approach or avoid novel objects [3]. Chicks of the domestic fowl, which belong to a precocial social species, appear to be endowed with predispositions to approach animate objects [2,22], given that in the absence of previous experience, young individuals prefer to approach facelike configurations [10], self-propelled objects [12], speed changes [30] and biologically-moving objects [11]. For young chicks, approaching choices are particularly important as they can influence imprinting. Filial imprinting is a process through which young chicks develop a strong social attachment, including following responses, to the first conspicuous objects they encounter in their life (for general reviews on chick's development and imprinting see Rogers [31], Bolhuis [32] and McCabe [26]). Although chicks can imprint on a variety of objects - including both natural and artificial objects -, specific colours, shapes, size and motion types induce stronger imprinting (see Introduction). Chicks' predispositions produce a bias in favour of naturalistic objects compared to artificial objects, as shown by the fact that once imprinted on a naturalistic object chicks cannot reverse their preference for an artificial object [33,34] or have a delayed reversal [35], although the opposite seems to be easier. Little is known though on the spontaneous preferences of chicks for approaching hollow or filled objects. This property can be particularly relevant to orient filial responses, because the presence of innards is associated with animate objects [23], that in the wild include social partners. Moreover, it has been observed that preschool children can reason about inside and outside features of objects [36], 14-month-old babies associate an object's behaviour more with internal than with external features [37], and preverbal infants (8-month-old) expect animate objects to possess insides [24]. In the case of human babies, spontaneous preferences in the absence of previous experience with hollow or filled objects can hardly be investigated. On the contrary, chicks are a convenient model as precocial and social species. We wondered whether the mere presence/absence of visible innards might trigger spontaneous approach preferences of young chicks for the first conspicuous objects encountered in their life, or whether experience might bias chicks preferences about the innards of social partners. To this aim we tested naïve and imprinted chicks using as hollow or filled objects orange cylinders of the size that can elicit filial responses.

In our experiments we consistently observed a preference of naïve chicks for approaching hollow objects. The same preference held for chicks that during imprinting had been exposed to objects occluded on their sides, that therefore were not explicitly filled or hollow. The preference for hollow objects decreased when chicks were imprinted for 24 hours on filled objects, suggesting that chicks are sensitive to this feature of the imprinting objects, and that even a brief experience can modify preferences for hollow/filled objects. Yet, we did not observe an increase of the preference for hollow objects after imprinting on hollow objects, and difference in performance between chicks imprinted on hollow and filled objects was not strong. This suggests that, after imprinting takes place, the feature of being hollow or filled is not crucial to change the perceived familiarity of the stimuli. Chicks imprinted on occluded cylinders that discover at test for the first time the hollow/filled distinction for the imprinting object approach more hollow objects, similarly to what naïve chicks do. This suggests that hollow stimuli - instead of opaque cylinders that could hide something potentially more interesting than an empty cavity - are more attractive for both naïve and imprinted chicks. To establish which property of hollow objects was attractive for chicks we ran a series of subsequent experiments to investigate whether chicks were attracted by hollow objects as hiding cavities, and/or whether the brightness and contrast of hollow objects were attractive cues that triggered exploration. Although inexperienced chicks spontaneously recognize the properties of occluding objects, and search objects behind barriers that completely occlude them [38], in our experiments chicks did not prefer larger hollow objects, in which they could more easily hide, to smaller hollow objects. This suggests that the preference for hollow objects is not mainly driven by the possibility to hide into them. On the contrary, chicks were more attracted by darker insides or darker “caps”. The attractive feature of hollow objects could be either the darker part inside the object (its shadows, which are a depth cue), or the higher contrast introduced by the presence of shadows. If the contrast but not the lower brightness was attracting the chicks, we expected them to have no preference when facing a choice between two scenes with the same (but opposite) contrast: a white disk on a black background and a black disk on a white background. Instead, in this setting chicks strongly preferred the black disk on a white background, suggesting that lower brightness of an object but not the contrast per se is attractive for chicks.

To sum up, naïve chicks exhibited a consistent preference for hollow objects, which was mainly mediated by the lower brightness of the insides, probably perceived as a depth cue. This preference could be modified by imprinting experience, by mere exposure of chicks to a filled object for 24 hours. At least for still objects such as the stimuli used in our experiments, the property of being “filled” does not make objects more attractive as imprinting objects for chicks of the domestic fowl. This suggests that cues possibily not connected to animacy might drive predisposed approach responses in chicks. Further experiments should clarify whether the preference for hollow *vs.* filled objects is modified introducing cues of animacy, such as the presence of movement or face configurations in the presented objects.

## Acknowledgements

GV was funded by the ERC Advanced Grant ERC-2011-ADG_20110406, Project No: 295517, PREMESOR

## Author contributions

Conceived and designed the experiments: EV, GV. Performed the experiments: JS, AMN, EV. Analyzed the data: EV, JS. Contributed materials/analysis tools: EV, JS. Drafted the manuscript: EV. Revised and approved the manuscript: EV, JS, AMN, GV.

